# Overlapping role of synaptophysin and synaptogyrin family proteins in determining the small size of synaptic vesicles

**DOI:** 10.1101/2024.05.29.596401

**Authors:** Daehun Park, Kenshiro Fujise, Yumei Wu, Rafael Luján, Sergio Del Olmo-Cabrera, John F. Wesseling, Pietro De Camilli

## Abstract

Members of the synaptophysin and synaptogyrin family are vesicle proteins with four transmembrane domains. In spite of their abundance in synaptic vesicle (SV) membranes, their role remains elusive and only mild defects at the cellular and organismal level are observed in mice lacking one or more family members. Here, we show that co-expression with synapsin of each of the four brain-enriched members of this family - synaptophysin, synaptoporin, synaptogyrin1 and synaptogyrin3 - in fibroblasts is sufficient to generate clusters of small vesicles in the same size range of SVs. Moreover, mice lacking all these four proteins have larger SVs. We conclude that synaptophysin and synaptogyrin family proteins play an overlapping function in the biogenesis of SVs and in determining their small size.

## Results and Discussion

A defining feature of synaptic vesicles (SVs) in nerve terminals is their very small size (1). However, the mechanisms accounting for such a characteristic remain poorly understood. We previously showed that ectopic expression in fibroblastic cells (COS7 cells) of synaptophysin, the second most abundant integral membrane protein of SVs (2), along with synapsin, a peripheral SV protein which has the property to assemble into a macromolecular condensate (3), is sufficient to generate liquid clusters of small exo-endocytic recycling vesicles similar to SVs of synapses in size and reminiscent of such vesicles in molecular composition (4, 5). We further showed that the cytosolic C-terminal tail of synaptophysin, which is negatively charged [pI = 3.91 (−4.1 at pH 7.3)] and harbors tyrosine containing repeats, mediates the interaction with the highly basic C-terminal tail of synapsin (4, 6). No such vesicle clusters were observed in fibroblasts when synapsin was expressed together with several other SV integral membrane proteins, including vesicle-associated membrane protein 2 (VAMP2), secretory carrier membrane 5 (SCAMP5), synaptotagmin 1, the vesicular glutamate transporter 1 (vGlut1) and the vesicular GABA transporter 1 (vGAT1), although these other proteins co-assembled with synaptophysin if co-expressed with synaptophysin and synapsin (5). These results suggested a specialized role of synaptophysin in the generation of vesicles with the size range of SVs which can be clustered by synapsin. However, no obvious morphological and functional changes at synapses can be observed in synaptophysin knock-out (KO) mice (7).

Mice express three other SV proteins that share structural similarities with synaptophysin and are believed to have a common evolutionary origin (8): synaptoporin (synaptophysin 2), synaptogyrin 1, and synaptogyrin 3 (Fig. 1A). Not only do these proteins have the same overall structure and topology as synaptophysin (Fig. 1A) - they are members of the tetraspan vesicle membrane protein (TVP) family with short cytosolic domains - but also share in their short C-terminal cytosolic tail features that were shown to be important for the interaction between synaptophysin and synapsin (4): highly negative charge [4.78 (−3 at pH 7.3) for synaptoporin; 3.25 (−7 at pH 7.3) for synaptogyrin 1; 4.25 (−3 at pH 7.3) for synaptogyrin 3] (Fig. 1A) and abundant presence of aromatic amino acids (6) (Fig. 1B). Interestingly, we have now found that in contrast to what was observed for other SV proteins tested besides synaptophysin (5), expression in COS7 cells of each of these three other proteins together with synapsin resulted in the formation of clusters of small vesicles similar in size to those generated by synaptophysin and synapsin co-expression (Fig. 1C and D). These results point to an overlapping role of these proteins in the biogenesis and clustering of SV.

**Fig. 1.**
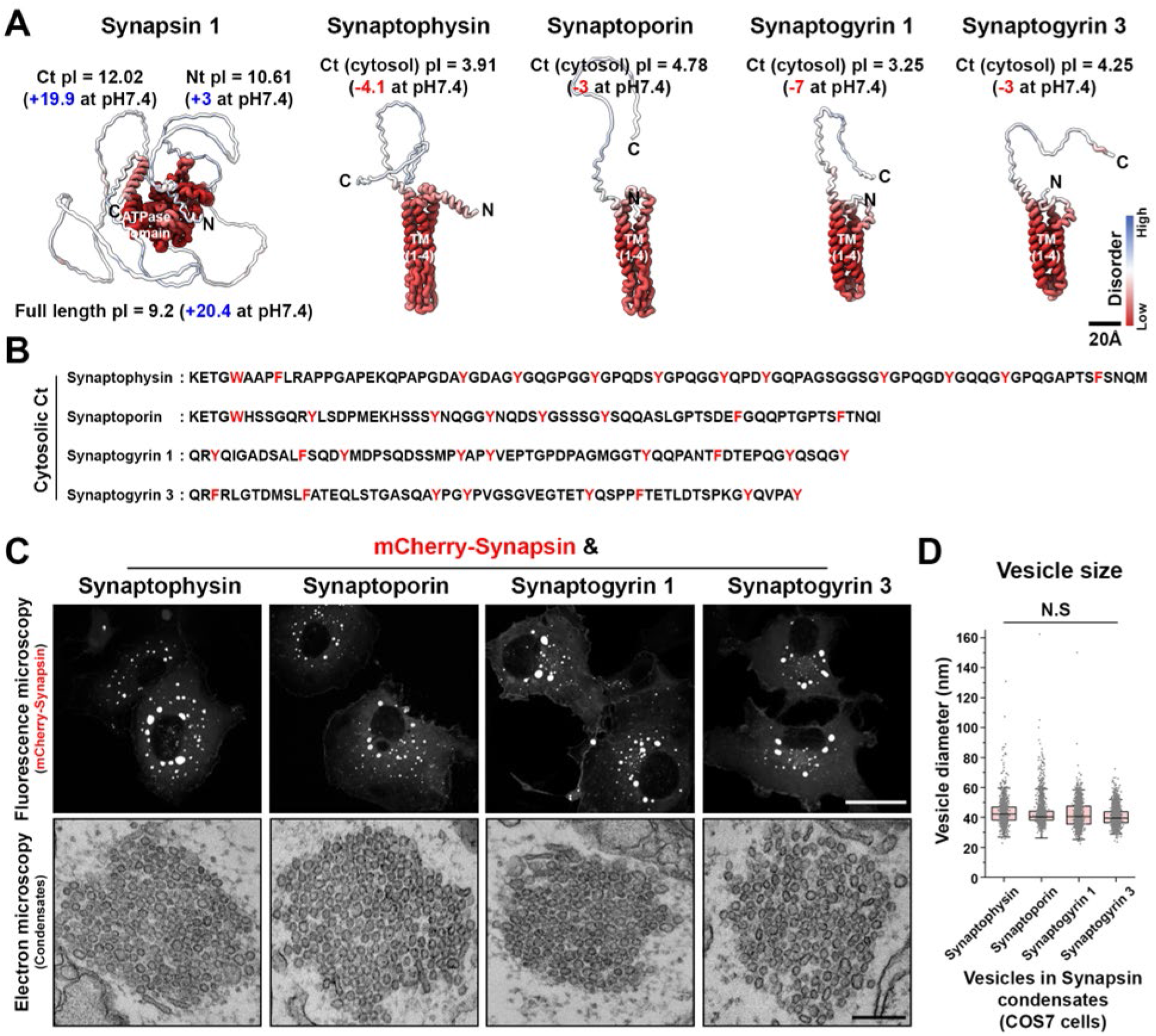
Synaptophysin and synaptogyrin family proteins form clusters of small vesicles with synapsin. *(A)* AlphaFold protein structures (human full length) and pI (isoelectric point) values of each protein. AFDB accession numbers: synapsin (AF-P17600-F1), synaptophysin (AF-P08247-F1), synaptoporin (AF-Q8TBG9-F1), synaptogyrin 1 (AF-O43759-F1), and synaptogyrin 3 (AF-O43761-F1). Colors represent disorderbility. *(B)* Cytosolic C-terminal sequence of each protein. *(C)* Representative confocal (top) and EM (bottom) images of condensates from COS7 cells expressing as indicated. *(D)* Size distribution of the vesicles. Box plots show the median line (midline), 25/75 percentiles (boxes), and 2SD (whiskers). 1469, 2109, 1978, and 2887 vesicles were measured for synaptophysin, synaptoporin, synaptogyrin 1, and synaptogyrin 3 expressing cells, respectively (from 4 independent experiments). N.S; no significant difference by one-way ANOVA with Tukey’s HSD post hoc test. Scale bars: *A* = 20 Å, *C* = 20 μm (top), 200 nm (bottom).

To validate this hypothesis, we examined by electron microscopy the morphology of synapses in stratum radiatum of the CA1 region of the hippocampus from a previously generated mouse model in which all four synaptophysin/synaptogyrin family proteins had been knocked-out [quadruple knockout (QKO)]. These mice are viable and fertile, revealing that the four tetraspanins are not essential for synaptic transmission (9). However, these mice are prone to seizures and studies of their synapses revealed increased neurotransmitter release in response to stimulation, consistent with subtle alterations in the neurotransmitter release machinery (9). Strikingly, we observed a clear increase in SV size relative to wild type [average SV diameter: 37.98 nm (WT) vs. 48.62 nm (QKO)] (Fig. 2A and B), consistent with a role of synaptophysin/synaptogyrin family members in determining the small size of SVs (Fig. 1C and D). Furthermore, spontaneous miniature synaptic currents (i.e., “minis) were also increased by about 28% in QKO CA1 pyramidal neurons (Fig. 2C-E), which matches the increase in mini size reported previously for calyx of Held synapses of QKO mice (9). Although, the increase in mini size did not scale linearly with the expected volume change (28% increase in diameter would imply a 110% increase in volume), it is consistent with the possibility that the individual vesicles contained more glutamate because of the larger size.

**Fig. 2.**
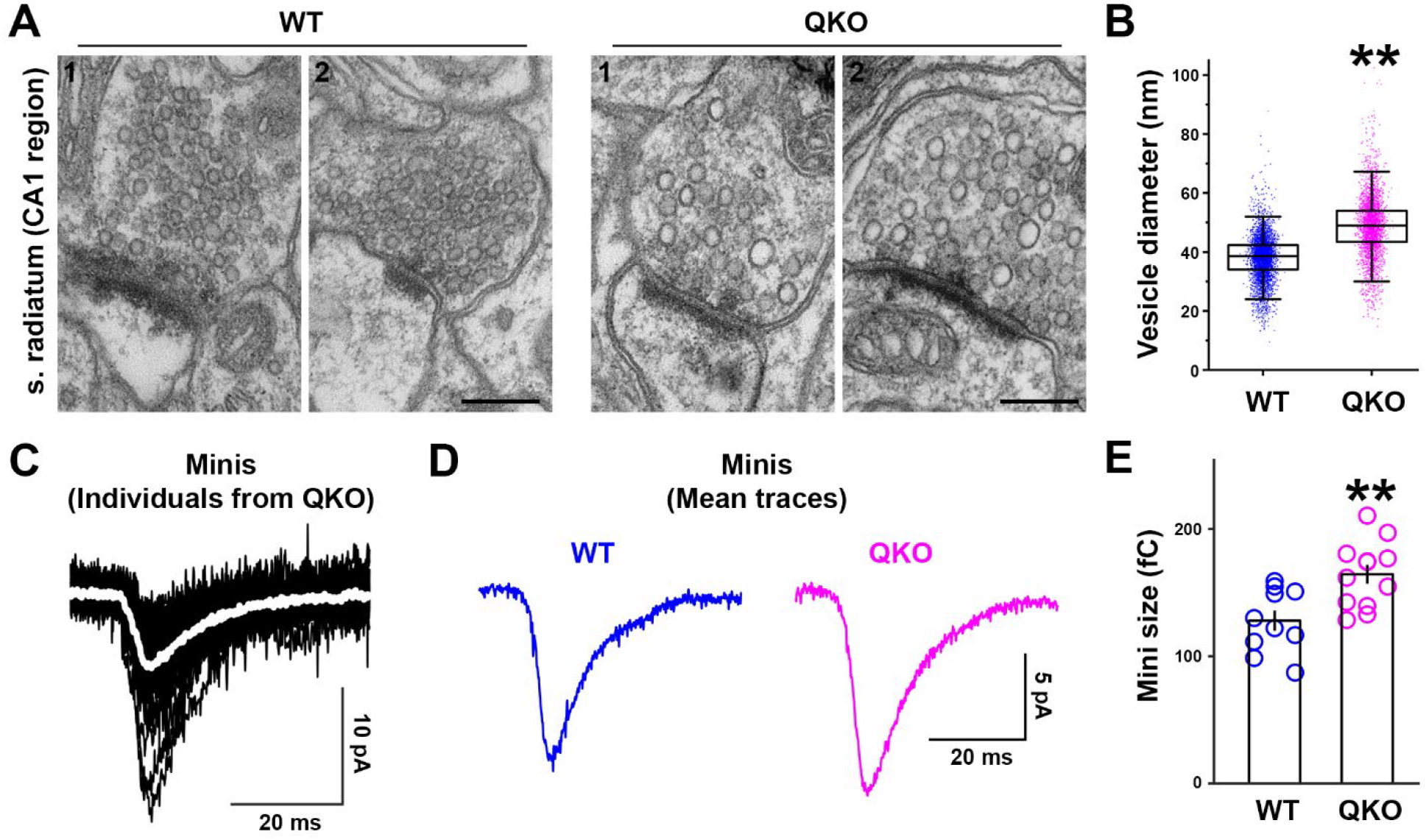
Increased size and quantal content of SVs in nerve terminals from synaptophysin, synaptoporin, synaptogyrin 1, and synaptogyrin 3 quadruple knock-out (QKO) mouse brains. *(A)* Representative EM micrographs from stratum radiatum in CA1 hippocampus of wild-type (WT) and QKO mutant mice. *(B)* Statistical analysis of *(A)*. 5691 (WT) and 4099 (QKO) vesicles were measured (from 4 different tissue sections from two animals for each group). \*\**p* < 0.01 by Student’s t-test. *(C-E)* Recordings were from CA1 pyramidal neurons, from 14- to 21-day-old mice. *(C)* Black traces are recordings of 32 minis from one QKO preparation. The superimposed white trace is the mean. *(D)* The mean of the means of all the minis detected for WT (blue) and QKO (magenta) neurons. *(E)* The current integrals of the mean mini for each neuron. \*\**p* < 0.01 by Student’s t-test. Scale bars: *A* = 200 nm.

We conclude that the four synaptophysin/synaptogyrin family proteins, while not required for the formation of neurotransmitter containing vesicles at synapses, may play a role in the acquisition of their characteristic very small size. Larger SVs had been observed at synapses of synaptogyrin null mutants of *Drosophila* that has a single synaptogyrin isoform and lacks a synaptophysin homolog (10). As we show here, synaptophysin/synaptogyrin family proteins are not only necessary to generate normal SVs but are also sufficient to generate small SV-like vesicles when expressed ectopically. A critical open question to be addressed is how these tetraspanins may determine vesicle shape and whether they do so by an intrinsic property or by recruiting to vesicles other factors with broad expression in neurons and non-neuronal cells.

## Materials and Methods

All animal experiments were approved by the corresponding Institutional Animal Care and Use Committee (IACUC of Yale University, and the Ethical Committee of the Generalitat Valenciana (2022/VSC/PEA/0255) for Alicante). Mouse colony maintenance, cell culture, transfection, imaging, and statistical analyses were done as previously described (4, 5, 9, 11). All experiments and analysis comparing wildtype and QKO were conducted blind to genotype. Additional details are provided in SI Appendix.

## Data Availability

All study data are included in the article and/or supporting information.

## Acknowledgements

This work was supported in part by grants from the National Institutes of Health of USA (NS036251 to P.D.C.), the Howard Hughes Medical Institute (P.D.C), the Agencia Estatal de Investigación of Spain (PID2021-125875OB-I00 to R.L. and PID2019-111131GB-I00 to J.F.W.), “ERDF A way of making Europe” (R.L.), the Junta de Comunidades de Castilla-La Mancha (SBPLY/21/180501/000064 to R.L.), the Generalitat Valenciana Prometeo Excellence Program (CIPROM2022/8 to J.F.W.), and the Brain Korea 21 Plus Program (D.P.).

## Author contributions

D.P., P.D.C. and J.F.W. designed the experiments. K.F. performed CLEM experiments. Y.W., and R.L. performed EM of mouse brain tissue. S.D.C. and J.F.W. performed electrophysiology. D.P. performed everything else. D.P. and P.D.C. wrote the paper. All authors read and approved the final manuscript.

