## Supplementary Information Appendix for "Overlapping role of synaptophysin and synaptogyrin family proteins in determining the small size of synaptic vesicles"

### Extended Materials and Methods

#### Cell culture and DNA transfection

COS7 cells were grown in DMEM (High Glucose) supplemented with 10% FBS and Penicillin-Streptomycin (100 U/ml) at 37 °C in 5% $CO_2$. For COS7 cell transfection, 60-80% confluent cells on 35 mm glass-bottom MatTek dish (P35G-1.5-14-CGRD) were transfected with total 0.5 µg of plasmid DNAs using Lipofectamine™ 2000 transfection reagent (Invitrogen). Cells were fixed and imaged after 24-48 hours of the transfection.

#### Correlative light and electron microscopy (CLEM) of cultured cells

COS7 cells were plated on 35 mm gridded grass-bottom dish and transfected as described above. Cells were fixed with 4% (v/v) PFA in Live Cell Imaging Buffer (Life Technologies) for 15min followed by three times washing with same buffer. Regions of interest were selected by fluorescence light microscopy imaging and their coordinates were identified using phase contrast. Imaging was performed with an Andor Dragonfly spinning-disk confocal imaging system with a Zyla CMOS camera and 60x plan apochromat objective (63x, 1.4 NA, oil). Cells were then processed as follows: Further fixation with 2.5% glutaraldehyde in 0.1 M sodium cacodylate buffer for 1 hour at room temperature; four 5 min washes with 0.1 M sodium cacodylate buffer; postfixation in 2% $OsO_4$ and 1.5% $K_4Fe(CN)_6$ (Sigma-Aldrich) in 0.1 M sodium cacodylate buffer on ice; four 5 min washes with Milli-Q water; staining with filtered 2% uranyl acetate dissolved in Milli-Q water for 1 hour; four 5 min washes with Milli-Q water; dehydration in a graded series of ethanol (50%, 75%, and 4 times in 100% for 5 min each); embedding in Embed 812. Ultrathin sections (40-60 nm) were observed in Talos L120C TEM microscope at 80 kV. Images were taken with Velox software and a 4k × 4 k Ceta CMOS Camera (Thermo Fischer Scientific). Except when noted, all EM reagents were from Electron Microscope Sciences. Quantification of vesicle diameters in COS7 cells was performed using randomly sampled images from clustered vesicles. Measurements were performed using FIJI (ImageJ).

### Electron microscopy of brain tissue

Mice were anesthetized and perfused with 2% paraformaldehyde and 2% glutaraldehyde in 0.1M PB (pH 7.4). Then, brains were dissected out and 60 μm coronal sections of brain regions containing the dorsal hippocampus were obtained using a Leica vibration microtome. After several washes in phosphate buffered saline, sections were postfixed with osmium tetroxide (1% in 0.1M PB) and *en bloc* stained with uranyl acetate (1% in distilled water). Sections were then dehydrated in ascending series of ethanol to 100% followed by propylene oxide and flat embedded on glass slides in Durcupan (Fluka, Barcelona, Spain). Ultrathin sections (40-60 nm) of the CA1 region of the hippocampus were observed either in a Philips CM10 TEM or in a Talos L120C TEM microscope at 80 kV. Images were taken with Velox software and a 4k × 4 k Ceta CMOS Camera (Thermo Fisher Scientific). Quantification of vesicle diameters at randomly selected synapses was performed using FIJI (ImageJ). The size of SVs in nerve terminals was measured under double-blind conditions.

### Electrophysiology

All experiments were conducted on 400 μm thick ex vivo transverse slices from the hippocampus with CA3 removed as described in (Wesseling and Lo 2002) except that slices were incubated at 34 °C for 45-60 min immediately after slicing and transferring to extracellular recording solution containing (in mM): 120 NaCl; 1.25 $NaH_2PO_4$; 26 $NaHCO_3$; 3.5 KCl; 10 glucose; 2 $MgCl_2$; 2 $CaCl_2$; 0.05 Picrotoxin, and 0.1 APV. Synaptic responses were recorded in whole cell voltage clamp mode by patch-clamping CA1 principle neurons with series resistance between 10 and 20 MΩ and data were only included for experiments where series resistance did not change by more than 20%. Minis were detected by hand in a blind fashion by presenting the evaluator with 200 ms-long fragments of electrophysiological traces, sampled at random from electrophysiological recordings from 22 neurons (22,113 fragments from 10 wildtype and 12 QKO recordings).

### Statistics

The Student's two-sample t-test was used to compare two independent groups. For multiple comparisons, ANOVA followed by Tukey's honest significant difference (HSD) post hoc test was applied. Data are presented as means ± SEM and the statistical significance was presented in the figure legends.
